# The role of hidden hearing loss in tinnitus: insights from early markers of peripheral hearing damage

**DOI:** 10.1101/2024.01.31.578195

**Authors:** Pauline Devolder, Hannah Keppler, Sarineh Keshishzadeh, Baziel Taghon, Ingeborg Dhooge, Sarah Verhulst

**Author notes:** Corresponding author, Technologiepark 126, 9050 Zwijnaarde, Belgium. Declarations of interest: none.

## Abstract

Since the presence of tinnitus is not always associated with audiometric hearing loss, it has been hypothesized that hidden hearing loss may act as a potential trigger for increased central gain along the neural pathway leading to tinnitus perception. In recent years, the study of hidden hearing loss has improved with the discovery of cochlear synaptopathy and several objective diagnostic markers. This study investigated three potential markers of peripheral hidden hearing loss in subjects with tinnitus: extended high-frequency audiometric thresholds, the auditory brainstem response, and the envelope following response. In addition, speech intelligibility was measured as a functional outcome measurement of hidden hearing loss. To account for age-related hidden hearing loss, participants were grouped according to age, presence of tinnitus, and audiometric thresholds. Group comparisons were conducted to differentiate between age- and tinnitus-related effects of hidden hearing loss. All three markers revealed age-related differences, whereas no differences were observed between the tinnitus and non-tinnitus groups. However, the older tinnitus group showed improved performance on low-pass filtered speech in noise tests compared to the older non-tinnitus group. These low-pass speech in noise scores were significantly correlated with tinnitus distress, as indicated using questionnaires, and could be related to the presence of hyperacusis. Based on our observations, cochlear synaptopathy does not appear to be the underlying cause of tinnitus. The improvement in low-pass speech-in-noise could be explained by enhanced temporal fine structure encoding or hyperacusis. Therefore, we recommend that future tinnitus research takes into account age-related factors, explores low-frequency encoding, and thoroughly assesses hyperacusis.

## 1 Introduction

Tinnitus is defined as the conscious awareness of a tonal or composite noise for which there is no identifiable corresponding external acoustic source (De Ridder et al., 2021) and affects 11.9% to 30.3% of the population (Hackenberg et al., 2023; Jarach et al., 2022; McCormack et al., 2016). Due to the heterogeneity of tinnitus, it remains challenging to identify its underlying mechanisms (Cederroth et al., 2019). Theories of tinnitus are often divided into bottom-up and top-down models, with bottom-up models explaining tinnitus based on peripheral hearing loss as a trigger and top-down models focusing on the maintenance of tinnitus perception at the cortical level. A combination of both is generally assumed, as originally proposed by Jastreboff (1990) in the ‘neurophysiological model’ and later extended in the ‘increased central noise model’ (Zeng, 2013) or the ‘interacting neural networks model’ (De Ridder et al., 2014), among other models. One bottom-up model is the central gain enhancement theory, which posits that increased neural activity through the auditory pathway compensates for reduced input due to peripheral hearing loss (Auerbach et al., 2014; Roberts & Salvi, 2019). In the majority of patients, peripheral hearing loss is visible on the audiogram. However, 8 to 10% of tinnitus patients have normal audiograms at conventional frequencies (Sanchez et al., 2005; Schaette & McAlpine, 2011). In such cases, other sensorineural deficits “hidden” by standard audiometry are considered a possible explanation for tinnitus generation.

A possible pathophysiological explanation for hidden hearing loss is cochlear synaptopathy (CS), i.e. damage to synapses between inner hair cells and afferent nerve fibers (ANFs) in the cochlea (Furman et al., 2013; Liberman & Kujawa, 2017). This phenomenon would be more likely to occur in synapses connected to nerve fibers with a low spontaneous spike rate (SR) (Furman et al., 2013). Compared to high-SR fibers, these have higher thresholds and a wider dynamic range, being sensitive to suprathreshold coding (Bharadwaj et al., 2014; Bourien et al., 2014; Furman et al., 2013). This suprathreshold coding is important for speech perception in noise and would be impaired in case of CS, while audiometric thresholds, determined by high-SR fibers, remain unaffected. Synaptopathy typically develops due to aging, after extensive noise exposure, or due to ototoxicity (Liberman & Kujawa, 2017; Sergeyenko et al., 2013). Since tinnitus is often reported in cases of age related, noise-induced or ototoxic hearing loss, with sometimes normal hearing thresholds, the link between tinnitus and cochlear synaptopathy is worth considering (Bhatt et al., 2016; Hackenberg et al., 2023; Knipper et al., 2013; Oosterloo et al., 2021).

In animals, CS can be determined post mortem by counting presynaptic vesicles and postsynaptic auditory nerve endings (Kujawa & Liberman, 2009). In humans, non-invasive electrophysiological measurements are used to determine synaptic loss. Several early markers of peripheral hearing loss have been proposed to map CS. A first marker concerns extended high frequency (EHF) audiometric thresholds, which are known to be increased due to age, noise exposure and ototoxicity, before affecting audiometric thresholds in the standard frequency range (Liberman et al., 2016; Mehrparvar et al., 2011). Secondly, based on the findings that CS would typically affect the temporal envelope (TENV) encoding of speech, the envelope following response (EFR), generated by amplitude modulated signals, is currently considered a promising objective electro-encephalography (EEG) test for hidden hearing loss (Bramhall, Beach, et al., 2019; Shaheen et al., 2015; Vasilkov et al., 2021; Verhulst et al., 2018). A last indicator that is considered as a potential marker for cochlear synaptopathy, is the wave-I amplitude of the auditory brainstem response (ABR) at suprathreshold level (Furman et al., 2013; Kujawa & Liberman, 2015; Schaette & McAlpine, 2011).

In addition to the ABR wave I, wave V is also used in both animal and human studies to demonstrate the central gain effect, which is assumed to be associated with tinnitus (Auerbach et al., 2014; Chen et al., 2021; Johannesen & Lopez-Poveda, 2021; Schaette & McAlpine, 2011). Human measurements indicate a significant reduction in the amplitude of wave I, but normal or even increased amplitudes of wave V in individuals with tinnitus and normal audiometric thresholds (Bramhall et al., 2018; Chen et al., 2021; Schaette & McAlpine, 2011). Correspondingly, animal studies of Auerbach et al. (2014) revealed decreased afferent input in the auditory nerve and cochlear nucleus, while, paradoxically, neural activity of the central auditory structures, including the inferior colliculus, the medial geniculate body, and the auditory cortex increased. For the measurement of central gain using the ABR in humans, ratios of wave V/I and wave I/V are commonly reported (Chen et al., 2021; Grose et al., 2019; Möhrle et al., 2016; Sergeyenko et al., 2013).

However, both CS and the central gain effect have also been observed as a result of aging (Chambers et al., 2016; Grose et al., 2019; Johannesen & Lopez-Poveda, 2021; Möhrle et al., 2016; Sergeyenko et al., 2013). Age-related effects were differentiated from tinnitus-effects in the ABR-study from Johannesen and Lopez-Poveda (2021), who concluded that the central gain effect is rather age-related and not associated with tinnitus. To unravel the influences of age and tinnitus, our study compared two young and two older groups, each with and without tinnitus. Our objective was to investigate the link between tinnitus and hidden hearing loss, especially CS, by considering three early markers of hidden peripheral hearing loss among individuals with normal hearing thresholds. Additionally, we assessed speech encoding as a more functional indicator of hidden hearing loss.

## 2 Materials and methods

### 2.1 Participants and audiometric thresholds

This study included four distinct test groups: young normal hearing without tinnitus (yNHnoT), young normal hearing with tinnitus (yNHT), older normal hearing without tinnitus (oNHnoT), and older normal hearing with tinnitus (oNHT) (see Table 1). All participants had (nearly) normal audiometric thresholds (see Figure 1), with a pure tone average (PTA; mean 1, 2, and 4 kHz) of maximum 20 dB HL. Frequency-specific thresholds were maximum 35 dB HL until 4 kHz and maximum 55 dB HL until 8 kHz. Despite slightly elevated thresholds in some older participants compared to the younger groups, we still refer to this group as normal hearing in this study, because their thresholds can be considered normal according to age (ISO 7029:2017(E)). However, this study primarily focuses on tinnitus-related differences within age groups to avoid confounding factors such as hidden hearing loss and cognitive decline associated with aging. Therefore, when interpreting differences between young and older normal-hearing groups, we consider these small audiometric differences. Hearing thresholds were measured using the modified Hughson-Westlake method (Carhart & Jerger, 1959). Participants were seated in a double-walled, sound-attenuated booth. Stimuli were presented using an Equinox Interacoustics audiometer on the conventional (half-)octave frequencies (0.125, 0.25, 0.5, 1, 2, 3, 4, 6 and 8 kHz) and EHFs of 10, 12.5, 14 and 16 kHz, through Interacoustics TDH-39 headphones and circumoral Sennheiser HAD-200 headphones respectively. Conductive hearing losses and middle/outer ear pathologies were excluded based on tympanometry and audiometric air-bone gaps when air conduction thresholds exceeded 20 dB HL. Bone conduction was measured using an Interacoustics bone vibrator placed on the mastoid. If the air-bone gap was 15 dB or more, participants were excluded from the study. For the rest of the protocol, only the best ear (based on overall thresholds) was measured. In the three subjects with unilateral tinnitus, the tinnitus ear was chosen.

**Figure 1:**
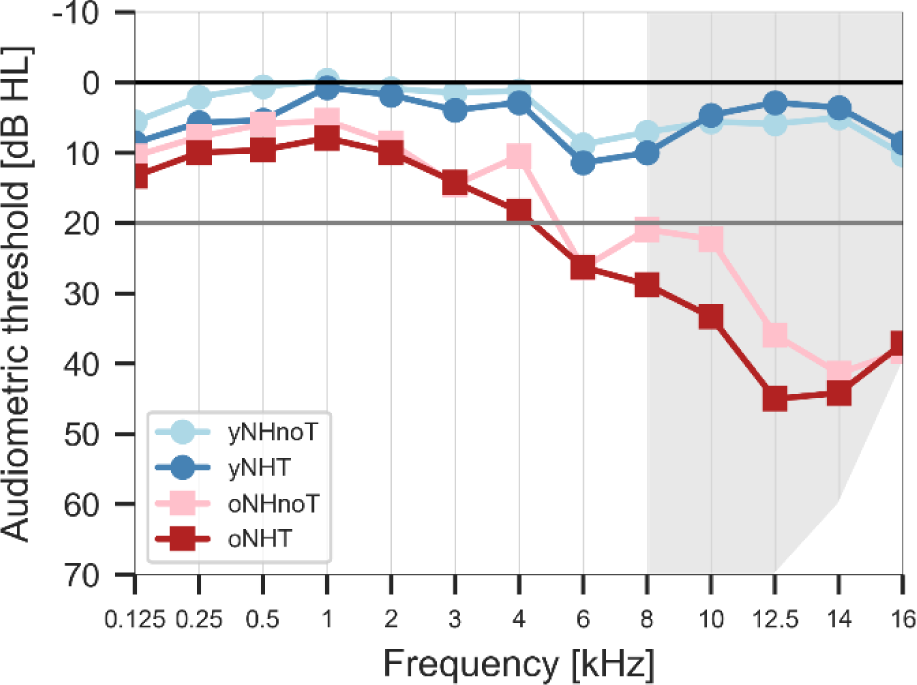
Averaged tonal audiograms per test group. The gray area indicates extended high frequencies, which were limited to 70 dB at 12.5 kHz, to 60 dB at 14 kHz and to 40 dB at 16 kHz. yNHnoT = young normal hearing without tinnitus; yNHT = young normal hearing with tinnitus; oNHnoT = older normal hearing without tinnitus; oNHT = older normal hearing with tinnitus.

**Table 1:** Mean demographic characteristics of the four test groups.

| Group | n | Age (years): min-max [m(SD)] | Test-ear: n(%) | PTA (dB HL): m(SD) |
| --- | --- | --- | --- | --- |
| yNHnoT | 17 (10 women, 7 men) | 20-24 [22.71 (0.849)] | 5 right (29.41%) | 0.59 (3.022) |
| yNHT | 14 (12 women, 2 men) | 20-25 [22.71 (1.437)] | 10 right (71.43%) | 1.78 (3.241) |
| oNHnoT | 11 (8 women, 3 men) | 40-55 [49.91 (5.127)] | 6 right (54.55%) | 8.18 (4.844) |
| oNHT | 12 (7 women, 5 men) | 41-55 [47.83 (6.235)] | 9 right (75.00%) | 12.08 (3.975) |

PTA = Pure Tone Average of 1, 2 and 4 kHz; yNHnoT = young normal hearing without tinnitus; yNHT = young normal hearing with tinnitus; oNHnoT = older normal hearing without tinnitus; oNHT = older normal hearing with tinnitus.

Exclusion criteria were a history of chronic otitis media, head trauma, ear surgery, retrocochlear lesions, endolymphatic hydrops, congenital ear malformations, neurological disorders and the use of ototoxic medication. The inclusion criterium for subjects with tinnitus was the presence of chronic tinnitus for at least 6 months. Participants were not allowed to be exposed to noise (concert, party, personal audio player) for at least 24 hours before participation.

All procedures complied with relevant laws and institutional guidelines and were approved by the ethical committee at Ghent University hospital (BC-00993-E05, 14/09/2020). Participants signed an informed consent before the experiment.

### 2.2 Questionnaires

Before the start of the experiment, all participants completed a questionnaire covering the following topics: (i) general sociodemographic questions, (ii) subjective hearing difficulties, (iii) tinnitus and (iv) noise exposure and use of hearing protection. Participants with tinnitus completed the Dutch version of following questionnaires assessing tinnitus and hyperacusis:

- The Tinnitus Sample Case History Questionnaire (TSCHQ) evaluates the history and characteristics of the tinnitus, including family history, onset, type of sound, laterality, loudness, pitch… (Langguth et al., 2007).
- The Tinnitus Handicap Inventory (THI) consists of 25 items divided into three subscales: functional (12 items), emotional (8 items), and catastrophic (5 items). Questions can be answered with ‘yes’ (4 points), ‘sometimes’ (2 points), or ‘no’ (0 points). The total score classifies individuals according to degrees of symptom severity: slight (0 to 16), mild (18 to 36), moderate (38 to 56), severe (58 to 76), and catastrophic (78 to 100) (Newman et al., 2008).
- The Tinnitus Functional Index (TFI) consists of 25 items divided into 8 subscales: intrusiveness, sense of control, cognitive interference, sleep disturbance, auditory issues, relaxation, quality of life, and emotional distress. Subjects responded using an 11-point visual analogue scale except for two questions rated between 0 and 100% that require a transformation from percentage to an 11-point scale. A total score of 26 or higher indicates that tinnitus has a significant impact on the person’s life. This questionnaire determines the severity of the tinnitus as well as the negative impact the person is experiencing (Meikle et al., 2012; Rabau et al., 2014).
- The Hyperacusis Questionnaire (HQ) consists of 14 questions that can be answered with ‘not true’, ‘sometimes true’, ‘often true’, ‘always true’ (4-point Likert scale). Hyperacusis is considered present in subjects with a score of 28 or more (Khalfa et al., 2002; Meeus et al., 2010).

### 2.3 Early markers of peripheral hearing damage

In addition to EHF audiometry, two EEG-based markers were measured. EFR and ABR stimuli were presented in alternating polarity and were delivered monaurally using an Intelligent Hearing Systems 2-channel opti-amp with ER-2 transducers. Electrodes were placed on Fz (non-inverting electrode (+)), on the ipsi and contralateral lobule (inverting electrodes (-)) and on Fpz near the nasion (ground electrode). Subjects were seated in a comfortable relax-chair while watching a muted video and were instructed to relax as much as possible, while remaining awake. Both ears were covered with earmuffs and all lights and nonessential electrical devices were turned off during the measurements.

#### 2.3.1 EFR

Since low-SR fibers are typically specialized in temporal envelope (TENV) encoding (Bharadwaj et al., 2014), the EFR focuses on the encoding of TENV using 110-Hz rectangular amplitude-modulated (RAM) pure tones as a stimulus (70 dB SPL, 4-kHz carrier, 500-ms epochs, 1000 repetitions, 100% modulated, 25 % duty cycle) (Van Der Biest et al., 2023; Vasilkov et al., 2021). The recordings were band-pass filtered using an 800th-order FIR filter (30-1500 Hz) in a zero-phase filtering procedure, epoched and baseline corrected. A bootstrapping approach was applied in the frequency domain to estimate the noise-floor and variability of the EFR (Keshishzadeh et al., 2020). EFR magnitudes were calculated as the sum of the signal-to-noise spectral magnitude at the fundamental frequency and its following three harmonics, i.e. 110, 220, 330 and 440 Hz (Van Der Biest et al., 2023; Vasilkov et al., 2021).

#### 2.3.2 ABR

ABRs were collected at 80 and 100 dB peSPL, using 11-Hz 80-μs clicks (4000 repetitions) and recordings were band-pass filtered (10-1500 Hz) using the same filter as for the EFR recordings, epoched and baseline corrected. The epochs were averaged to obtain the ABR waveform in which waves I, III, V and VI were manually peak-picked by trained audiologists to identify the respective ABR peak-to-baseline amplitudes.

### 2.4 Speech (in noise) test

Based on the physiological evidence discussed by Verschooten et al. (2019) and the binaural human phase-locking limit near 1.5 kHz (Brughera et al., 2013), we assume that human TENV coding dominates over temporal fine structure (TFS) coding for high frequencies. With this assumption, we investigated speech intelligibility using low-frequency (<1.5 kHz) and high-frequency (>1.65 kHz) filtered speech, which would provide either predominant TFS or TENV cues respectively. Speech intelligibility was quantified using the speech reception threshold (SRT) using the Apex-3 test platform (Francart et al., 2008). The tests were performed monaurally (to the ear with the best audiometric thresholds) with Sennheiser HD-300 headphones and a Fireface UCX soundcard. The Flemish 5-word Matrix sentence test (Luts et al., 2014) was conducted in an adaptive tracking procedure to determine the SRT. The following conditions were tested with and without stationary speech-shaped background noise of 70 dB SPL: broadband (BB) filtered, low-pass (LP) filtered (1.5-kHz 1024th-order FIR filter) and high-pass (HP) filtered (1.65-kHz 1024th-order FIR filter).

### 2.5 Statistics

EEG data was analyzed using MATLAB and statistical analyses were performed with Python and IBM SPSS Statistics 29. A two-way analysis of variance (ANOVA) was conducted to investigate the effects of two independent variables (tinnitus presence and age group). The significance level for all analyses was set at α = 0.05. To further explore significant effects identified by the ANOVA, group comparisons were conducted using two-sided independent Student’s t-tests or Mann-Whitney U tests. Depending on the normality of the variables (using the Shapiro Wilk test), parametric or nonparametric group statistics were performed. Since four comparisons were included in the group comparisons, a Bonferroni correction was applied, bringing the significance level to α = 0.0125:

- yNHnoT with yNHT to verify the tinnitus-related effect in the younger group.
- oNHnoT with oNHT to verify the tinnitus-related effect in the older group.
- yNHnoT with oNHnoT to verify any age-related effect.
- yNHT with oNHT to verify any age-related effect.

For all significant comparisons, the Cohen’s d effect size is reported, with d ≥ 0.800 interpreted as a large effect.

To further explore linear relationships between continuous variables, the Pearson or Spearman correlation coefficient was calculated (depending on their normality), and corresponding p-values were reported with a significance level of α=0.05. During correlation analyses with LP SPIN scores, additional Bonferroni corrections were applied to address multiple comparisons. 21 comparisons were performed in total, but were not independent of each other. Since THI scores highly correlate with subscale scores (functional: *r* = 0.948; emotional: *r* = 0.932; and catastrophic: *r* = 0.827), we considered 18 independent comparisons, reducing the significance level to α = 0.0028.

## 3 Results

### 3.1 Tinnitus questionnaires

Results of the main tinnitus-related questionnaire items are summarized in Table 2. Tinnitus duration was significantly shorter in the young tinnitus group (yNHT) compared to the older group (oNHT) (*t*(21) = -2.837; *p* = 0.01), while tinnitus loudness did not differ significantly (*p* > 0.05). Analysis of the THI and TFI scores revealed no significant differences in tinnitus distress between the two groups, even for their separate subcategories (*p* > 0.05). In addition, hyperacusis scores based on the HQ questionnaire did not differ significantly between our two groups (*p* > 0.05).

**Table 2:**
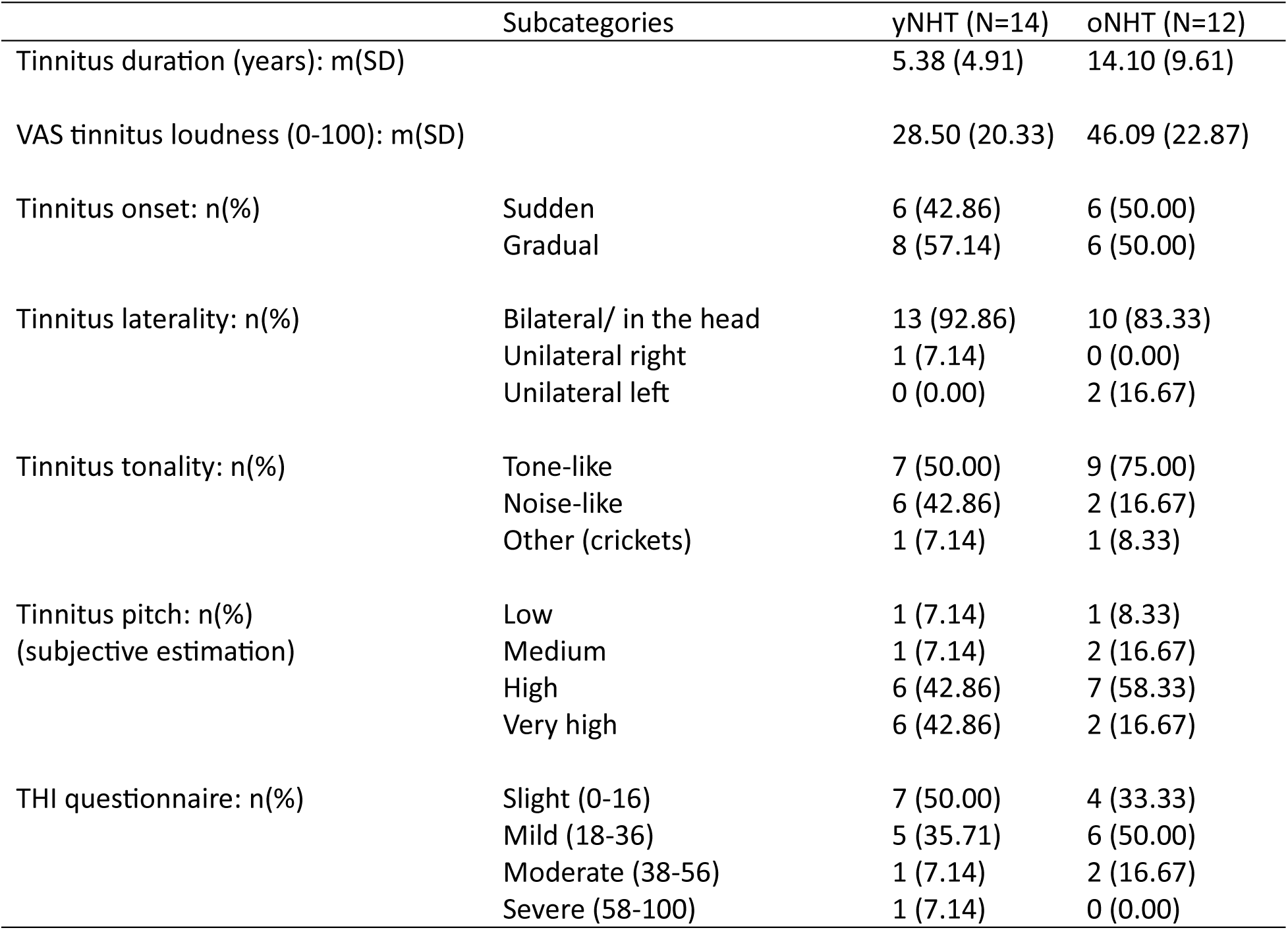
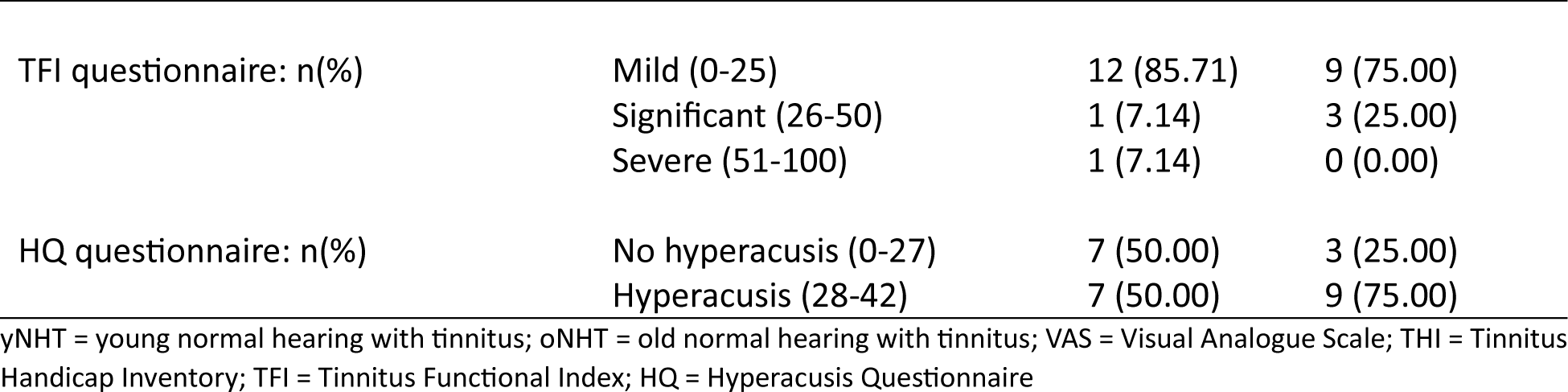
Tinnitus characteristics and questionnaire results of the two groups with tinnitus.

### 3.2 Early markers of peripheral hearing damage

#### 3.2.1 EHF audiometry

The marker considered first is the mean of hearing thresholds across EHFs. Significant age-related differences were observed (Table 3), with better thresholds for the young group. Group comparisons showed that his age effect was present within both the non-tinnitus (*t*(26) = -6.52; *p* < 0.0001) and tinnitus groups (*t*(24) = -7.23; *p* < 0.0001) (see Figure 2c). Nevertheless, no tinnitus-related effects were found (p > 0.05; Table 3). Standard audiometric thresholds were split up in low and high frequencies. When considering the mean of the low frequencies (125 to 1000 Hz), both tinnitus presence and age show significant differences in the two-way ANOVA analysis (Table 3). Group comparison only showed tinnitus-related differences between the young groups (*t*(29) = -2.63; *p* = 0.0136), which disappears after Bonferroni correction (p > 0.0125). Age-related differences were still present after Bonferroni correction in both non-tinnitus and tinnitus groups (Figure 2a; yNHnoT and oNHnoT: *t*(26) = -3.78; *p* = 0.0008; yNHT and oNHT: *t*(24) = -3.30; *p* = 0.0030). The mean of high frequency thresholds only revealed age-related differences, in both tinnitus groups (Table 3; Figure 2b; yNHnoT and oNHnoT: *t*(26) = -5.54; *p* < 0.0001; yNHT and oNHT: *t*(24) = -6.38; *p* < 0.0001). Effect sizes of these significant comparisons were large, ranging between d = 1.462 and d = 2.915.

**Figure 2:**
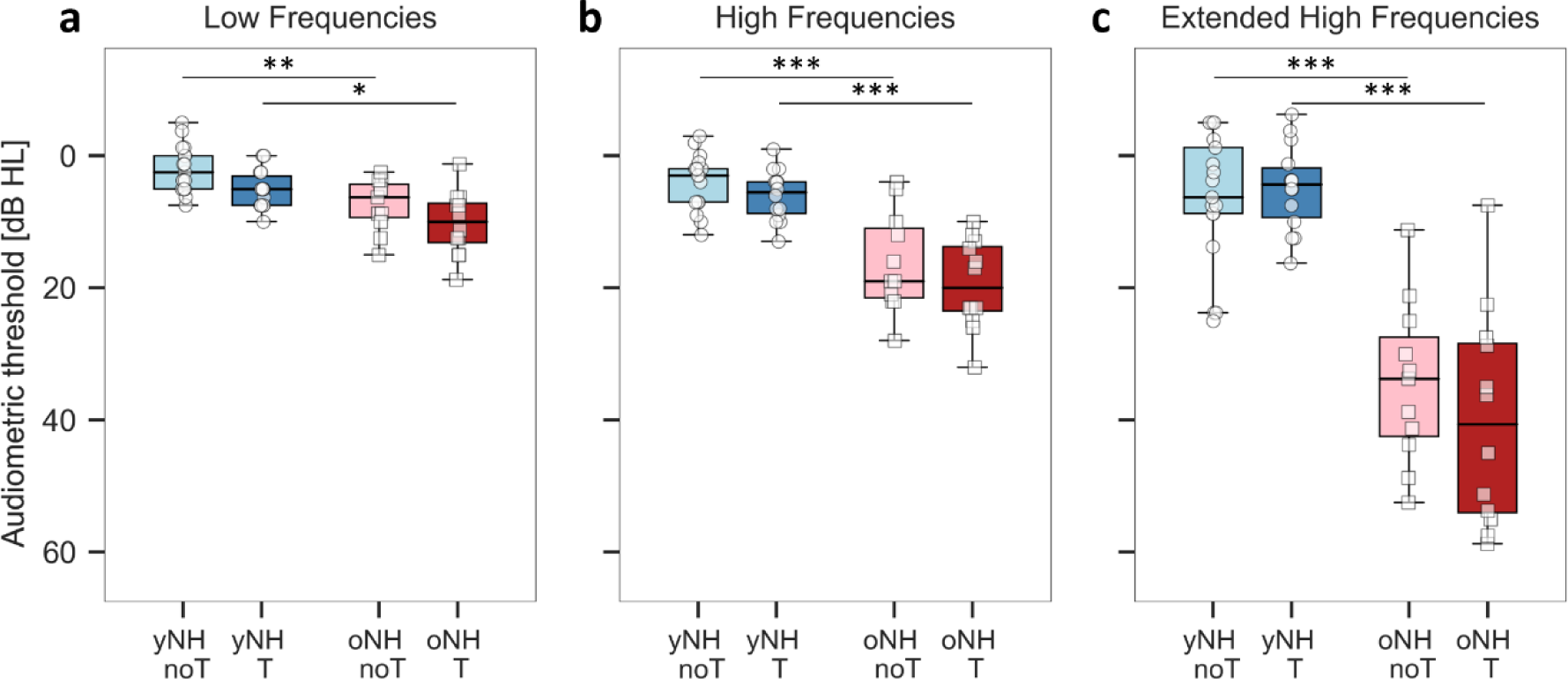
Boxplots of (a) the mean of the low frequent audiometric thresholds (125, 250, 500 and 1000 Hz), (b) the mean of the high frequent audiometric thresholds (2000, 3000, 4000, 6000 and 8000 Hz) and (c) the mean of the extended high frequency (EHF) audiometric thresholds (10, 12.5, 14 and 16 kHz) for each test group. yNHnoT = young normal hearing without tinnitus; yNHT = young normal hearing with tinnitus; oNHnoT = older normal hearing without tinnitus; oNHT = older normal hearing with tinnitus. *p ≤ 0.0125; **p ≤ 0.001; ***p ≤ 0.0001.

**Table 3:** Outcomes of two-way ANOVA analyses for each parameter, considering tinnitus presence, age group and the interaction effect between tinnitus presence and age group.

|  | Presence<br>of tinnitus | Age group | Interaction |
| --- | --- | --- | --- |
| LF audiometry (125, 250, 500, 1000 Hz) | **p = 0.007**<br>F(1, 50) = 7.926 | **p < 0.001**<br>F(1, 50) = 24.979 | p = 0.894 |
| HF audiometry (2, 3, 4, 6, 8 kHz) | p = 0.083 | **p < 0.001**<br>F(1, 50) = 70.497 | p = 0.698 |
| EHF audiometry (10, 12.5, 14, 16 kHz) | p = 0.562 | **p < 0.001**<br>F(1, 50) = 98.580 | p = 0.257 |
| EFR 4000 Hz | p = 0.983 | **p < 0.001**<br>F(1, 48) = 43.728 | p = 0.939 |
| ABR 80 dB peSPL, amplitude wave I | p = 0.279 | **p < 0.001**<br>F(1, 49) = 21.881 | p = 0.907 |
| ABR 80 dB peSPL, amplitude wave V | p = 0.769 | **p < 0.001**<br>F(1, 50) = 16.263 | p = 0.644 |
| ABR 80 dB peSPL, amplitude wave I/V | p = 0.684 | **p < 0.001**<br>F(1, 49) = 12.412 | p = 0.846 |
| ABR 80 dB peSPL, latency wave I | p = 0.870 | **p = 0.046**<br>F(1, 49) = 4.178 | p = 0.764 |
| ABR 80 dB peSPL, latency wave V | p = 0.879 | **p = 0.003**<br>F(1, 50) = 9.787 | p = 0.969 |
| ABR 100 dB peSPL, amplitude wave I | p = 0.821 | **p = 0.010**<br>F(1, 49) = 7.139 | p = 0.683 |
| ABR 100 dB peSPL, amplitude wave V | p = 0.298 | **p = 0.001**<br>F(1, 49) = 11.885 | p = 0.956 |
| ABR 100 dB peSPL, amplitude wave I/V | p = 0.280 | p = 0.083 | p = 0.207 |
| ABR 100 dB peSPL, latency wave I | p = 0.362 | p = 0.735 | p = 0.294 |
| ABR 100 dB peSPL, latency wave V | p = 0.376 | **p = 0.013**<br>F(1, 49) = 6.691 | p = 0.314 |
| Speech in quiet low pass | p = 0.180 | **p = 0.041**<br>F(1, 50) = 0.765 | p = 0.827 |
| Speech in quiet high pass | p = 0.159 | **p < 0.001**<br>F(1, 49) = 34.424 | p = 0.749 |
| Speech in noise low pass | **p = 0.011**<br>F(1, 50) = 6.945 | p = 0.208 | p = 0.107 |
| Speech in noise broad band | p = 0.185 | **p = 0.006**<br>F(1, 50) = 8.083 | p = 0.761 |
| Speech in noise high pass | p = 0.282 | **p < 0.001**<br>F(1, 50) = 25.071 | p = 0.798 |

EFR = Envelope Following Response; ABR = Auditory Brainstem Response, LF = Low Frequency, HF = High frequency, EHF = Extended High Frequency

#### 3.2.2 EFR

Similar to the EHF audiometry findings, only age-related effects were observed for EFR magnitudes in both tinnitus and non-tinnitus groups (Table 3; Figure 3). The yNHnoT group showed significantly better EFR-magnitudes compared to the oNHnoT group (*t*(26) = 4.76; *p* = 0.0001; d = 1.842) and the yNHT group scored significantly better than oNHT (*t*(22) = 4.60; *p* = 0.0001; d = 1.884). No tinnitus-related differences were found (*p* > 0.05).

**Figure 3:**
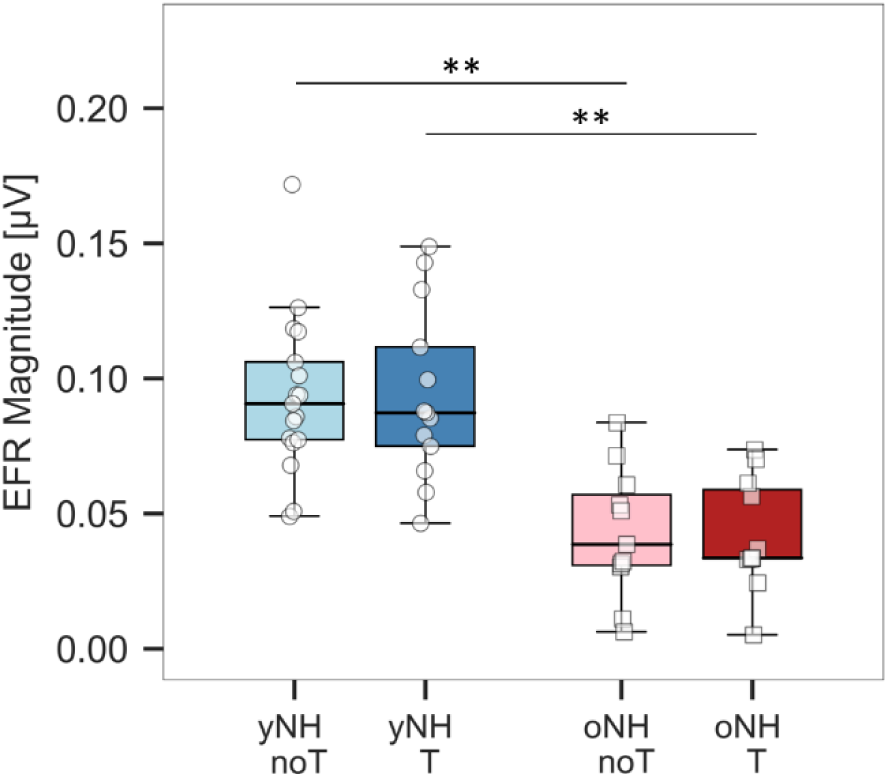
Boxplots of the envelope following response at 4000 Hz for each test group. yNHnoT = young normal hearing without tinnitus; yNHT = young normal hearing with tinnitus; oNHnoT = older normal hearing without tinnitus; oNHT = older normal hearing with tinnitus. *p ≤ 0.0125; **p ≤ 0.001; ***p ≤ 0.0001.

#### 3.2.3 ABR

The average ABR peaks and troughs at 80 dB peSPL are visualized in Figure 4a. Differences in amplitudes and latencies of peak wave I and V were analyzed at both 80 and 100 dB peSPL. At 80 dB peSPL, age-related amplitude decreases were observed for wave I (Table 3), this effect was present in the non-tinnitus groups (*t*(25) = 2.71; *p* = 0.0121), as well as in the tinnitus groups (*t*(24) = 4.97; *p* < 0.0001) (see Figure 4b). Confirming the previous discussed markers, no significant tinnitus-related differences were detected (*p* > 0.05). Wave-I latencies at 80 dB peSPL showed age-related effects in the two-way ANOVA analysis (Table 3), but not in the comparisons between the four groups (p > 0.05). At 100 dB peSPL, wave I showed age-related amplitude-differences (Table 3), not persisting in the group comparisons (p > 0.05). No wave-I latency differences were observed (*p* > 0.05).

**Figure 4:**
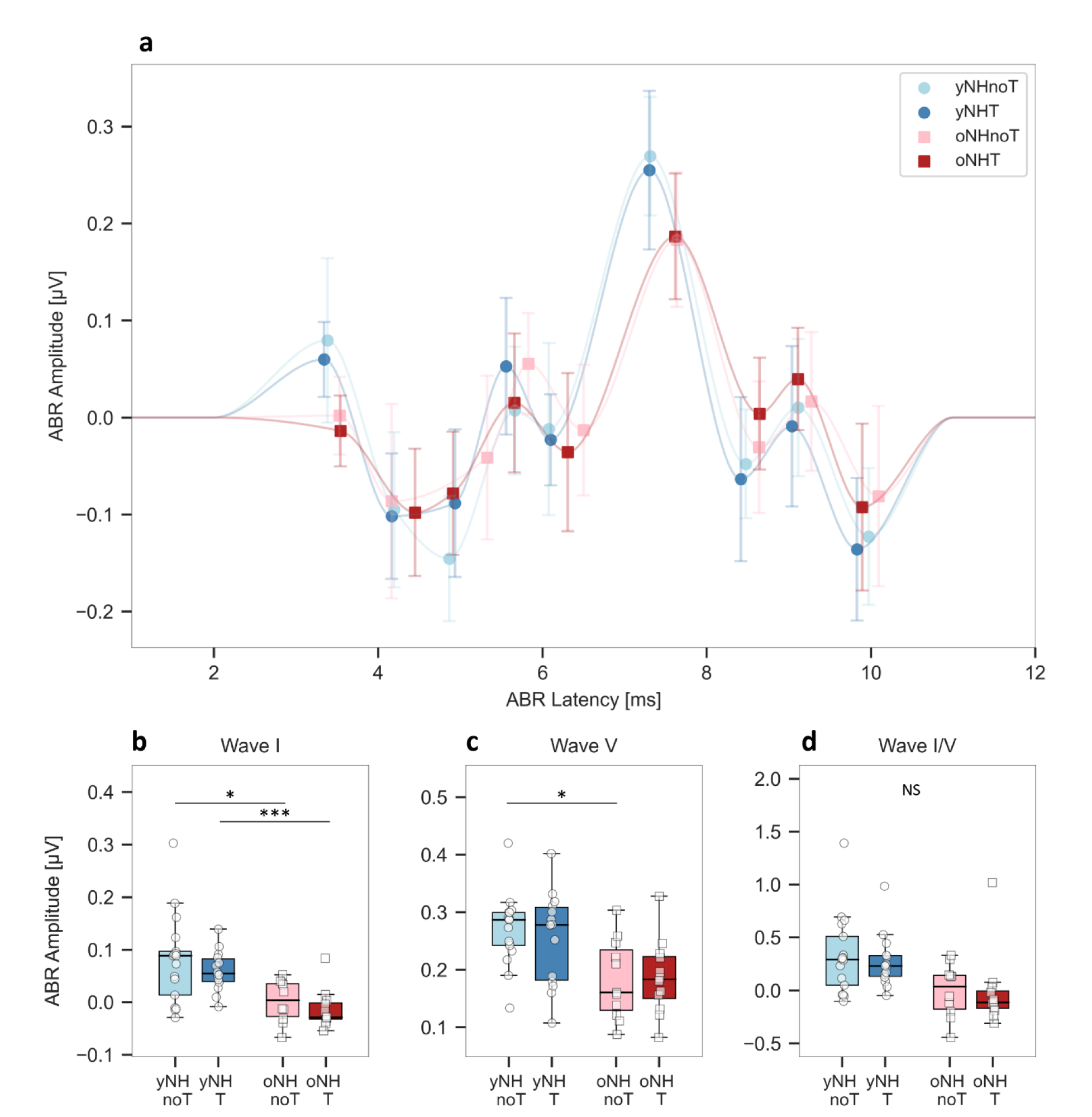
Auditory brainstem responses (ABR) with an 80 dB peSPL click. (a) Mean ABR-latency and -amplitude values for each peak-picked peak and trough (P1, N1, N2, P3, N3, P5, N5, P6, N6). Means are presented for each test group, and connected with light, curved trend lines to improve clarity. (b) Boxplot of the positive wave-I (P1) amplitudes for each test group. (c) Boxplot of the positive wave-V (P5) amplitudes for each test group. (c) Boxplot of the wave-I/V ratio for each test group. yNHnoT = young normal hearing without tinnitus; yNHT = young normal hearing with tinnitus; oNHnoT = older normal hearing without tinnitus; oNHT = older normal hearing with tinnitus. *p ≤ 0.0125; **p ≤ 0.001; ***p ≤ 0.0001; NS = not significant

The wave-V amplitude at 80 dB peSPL was significantly different for age (Table 3), only for the non-tinnitus groups after Bonferroni correction (*t*(26) = 3.46; *p* = 0.0019) (Figure 4c). At 100 dB peSPL, this age-related effect also occurred, but was not significant after Bonferroni correction in the group comparisons (p > 0.0125). Wave V also showed significant latency differences related to age (Table 3), with the young tinnitus group showing shorter latencies at 80 dB peSPL than the older group (*t*(24) = -3.06; *p* = 0.0053). At 100 dB peSPL the latency differences (Table 3) did not persist in the group comparisons after Bonferroni correction (*p* > 0.0125). Consistent with the previous analyses, no tinnitus-related differences were observed for wave V at either intensity level (*p* > 0.05).

In Figure 4d, the ABR wave-I/V ratio was compared between the test groups, with lower values representing more central gain. Only at 80 dB peSPL, differences were found between the age groups (Table 3). Group comparisons revealed these differences both within the tinnitus and non-tinnitus groups, but did not persist after Bonferroni correction (p > 0.0125). Again, no tinnitus-related differences appeared (*p* > 0.05). Effect sizes of ABR-related significant differences were large, ranging between d = 1.078 to d =1.957.

### 3.3 Speech encoding

Speech perception in quiet (SPIQ) was evaluated in a LP- and HP-filtered condition. Both conditions showed age-related differences (Table 3). No group differences were found in the LP condition (p > 0.05; Figure 5a), but significant differences were observed in the HP condition (Figure 5b), with young participants performing better than the older ones in both the tinnitus (*t*(26) = -4.33; *p* = 0.0002) and non-tinnitus groups (*t*(23) = -3.98; *p* = 0.0006). These HP SPIQ scores correlated strongly to the mean audiometric thresholds at high frequencies over all groups (2000-8000 Hz; *ρ* = 0.8019; *p* < 0.0001).

**Figure 5:**
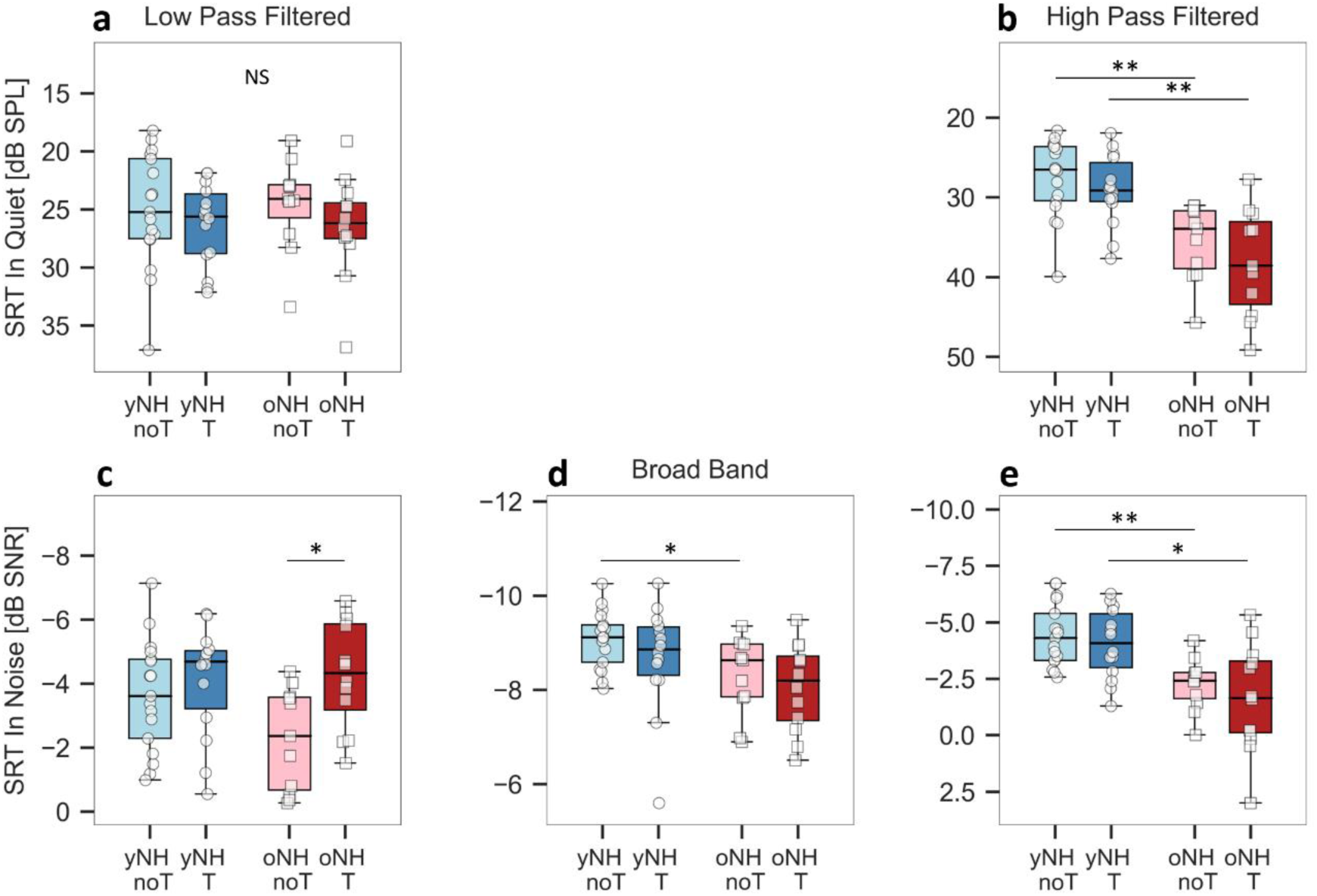
Boxplot of (a) the low-pass (LP) filtered speech in quiet test; (b) the high pass (HP) filtered speech in quiet test; (c) the LP filtered speech in noise test; (d) the non-filtered (BB) speech in noise test; (e) the HP filtered speech in noise test; all compared between test groups. SRT = Speech Reception Threshold; yNHnoT = young normal hearing without tinnitus; yNHT = young normal hearing with tinnitus; oNHnoT = older normal hearing without tinnitus; oNHT = older normal hearing with tinnitus. *p ≤ 0.0125; **p ≤ 0.001; ***p ≤ 0.0001; NS = not significant.

Speech perception in noise (SPIN) was evaluated in three conditions: LP, BB and HP. Again, age was found to have a significant impact on performance (Table 3), with the young groups scoring better than the older groups in the HP condition for both non-tinnitus (*t*(26) = -4.36; *p* = 0.0002) and tinnitus groups (*t*(24) = - 3.09; *p* = 0.0050) (Figure 5e). In the BB condition, young participants also scored better, but only in the non-tinnitus groups (*t(*26) = -2.84; *p* = 0.0086) (Figure 5d). Surprisingly, a tinnitus-related difference surfaced in the LP condition (Table 3), with older participants with tinnitus (oNHT) scoring significantly better than the older without tinnitus (oNHnoT) (*t*(21) = 2.91; *p* = 0.0084), and performing equally well as the young groups (*p* > 0.05) (Figure 5c). This effect was not observed in the young group (p > 0.05).

Since the observations of improved speech-in-noise understanding in tinnitus patients are rather unexpected, we explored several correlations between LP SPIN scores and the other parameters measured in this study. While HP SPIN correlated with mean high-frequency audiometric thresholds (*ρ* = 0.5460; *p* < 0.0001), the low-pass condition did not correlate with the mean low-frequency audiometric thresholds, nor did it correlate with LP SPIQ, age, EHF audiometry, ABR amplitudes at 80 and 100 dB peSPL or EFR magnitudes at 4000Hz (*p* > 0.05). Again, significant comparisons showed large Cohen’s d effect sizes (d = 1.101 to d = 1.674).

### 3.4 Link between speech-in-noise intelligibility and tinnitus distress

To further explore possible factors related to LP SPIN scores, questionnaires outcomes were considered using correlations analyses.

#### 3.4.1 Tinnitus Handicap Inventory (THI)

The THI questionnaire, assessing tinnitus distress, revealed significant correlations within the older group (Figure 6a). Higher THI scores correlated with better LP SPIN (*r* = -0.6862; *p* = 0.0137; 95% CI [-0.904, - 0.185]), a trend not observed in the younger group (*p* > 0.05). Further exploration into THI subscales highlighted significant correlations in the older tinnitus group for the functional (*r* = -0.6715; *p* = 0.0168) and emotional subscale (*r* = -0.7429; *p* = 0.0056), but not for catastrophic subscale (*p* > 0.05) scores. No correlations were found between THI scores and other speech conditions, nor for the subscales (*p* > 0.05). Due to the large number of correlation analyses already performed, we implemented a Bonferroni correction, bringing the significance level to 0.0028. After conducting this more strict analysis, the correlation with THI is no longer statistically significant.

**Figure 6:**
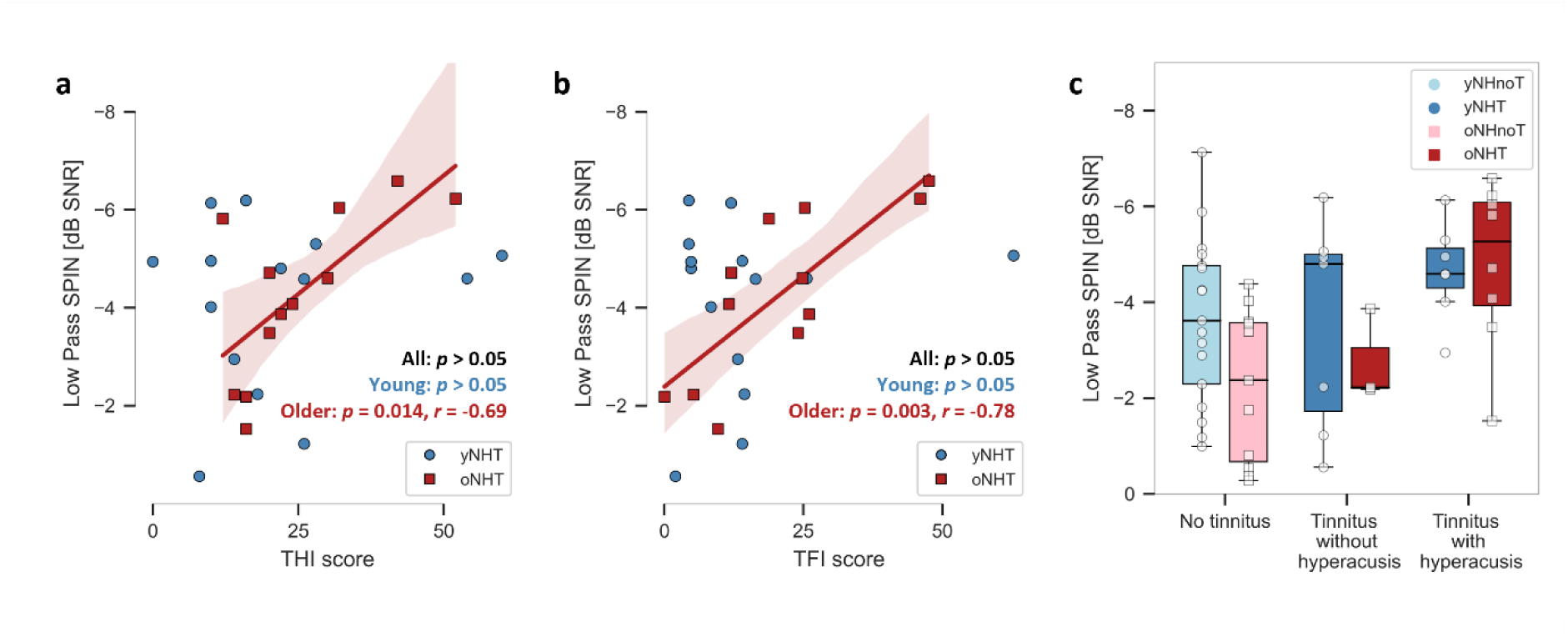
Scatter plots of the relation between low-pass filtered speech-in-noise (SPIN) scores and (a) Tinnitus Handicap Inventory (THI) scores and (b) Tinnitus Functional Index (TFI) scores. (c) Boxplots of low-pass-filtered SPIN scores for three groups: individuals without tinnitus, those with tinnitus but without hyperacusis (scores < 28 on the hyperacusis questionnaire), and those with tinnitus and hyperacusis (scores ≥ 28 on the hyperacusis questionnaire). yNHnoT = young normal hearing without tinnitus; yNHT = young normal hearing with tinnitus; oNHnoT = older normal hearing without tinnitus; oNHT = older normal hearing with tinnitus.

#### 3.4.2 Tinnitus Functional Index (TFI)

Additionally, TFI scores also represent tinnitus distress and showed a significant correlation with LP SPIN in the older group (*r* = -0.7843; *p* = 0.0025; 95% CI [-0.937 , -0.383]; see Figure 6b). When splitting up in subscales, this link was especially present for the relaxation (*ρ* = -0.8035; *p* = 0.0016) and emotional subscale (*ρ* = -0.5937; *p* = 0.0418), with the latter also correlating in the young group (*ρ* = -0.6056; *p* = 0.0217). The other subscales showed similar trends without reaching statistical significance (*p* > 0.05). After Bonferroni correction, two correlations remained, namely the total TFI score and the relaxation subscale. No correlations were observed between TFI scores and other speech conditions, nor for the subscales (*p* > 0.05).

#### 3.4.3 Hyperacusis Questionnaire (HQ)

No significant correlations were found between HQ scores and LP SPIN (*p* > 0.005). Based on the cutoff of 28, the group was split into tinnitus patients with and without hyperacusis and visually compared with the non-tinnitus group scores in Figure 6c. Group statistics were not performed due to the small groups sizes when splitting up for hyperacusis. Nevertheless, notable trends emerged since older participants with both tinnitus and hyperacusis appear to score better on LP SPIN than both groups without hyperacusis. Similarly, in the young groups, improved scores were visually observed for the hyperacusis group, despite the wide range of LP SPIN scores in the non-tinnitus group. Additionally, only in the young group, HQ scores were correlated with BB SPIN (*r* = -0.7582; *p* = 0.0017). No other correlations between HQ scores and other speech conditions were observed (*p* > 0.05).

## 4 Discussion

The contribution of peripheral hidden hearing loss in the pathogenesis of tinnitus remains unclear. Generally, hidden hearing loss was considered as a plausible explanation, especially in individuals with normal hearing thresholds. This study explored the occurrence of hidden hearing loss in tinnitus patients by measuring three potential markers: EHF audiometry, EFR, and ABR wave I. Additionally, we investigated the phenomenon of enhanced central gain through the ABR wave-I/V ratio and assessed speech intelligibility as a functional indicator of hidden hearing loss. Normal hearing groups with and without tinnitus were subdivided in young and older participants in order to consider age-related effects of hidden hearing loss.

### 4.1 Cochlear synaptopathy does not seem to be the driving force behind tinnitus

By subdividing the groups by age, this study revealed no statistically significant tinnitus-related differences in any markers associated with peripheral hidden hearing loss. However, significant differences were observed between the younger and older groups for all markers, which could be explained by age-related effects such as hidden hearing loss or higher hearing thresholds. CS could be an explaining factor, since EFR-magnitudes were smaller in the older groups (Vasilkov et al., 2021). Nevertheless, the absence of significant differences between tinnitus and non-tinnitus groups suggests that tinnitus is not associated with hidden hearing loss or CS. Rather, reduced CS-markers seem to be associated with age-related factors. This is consistent with the findings of Johannesen and Lopez-Poveda (2021), who attributed differences in ABR amplitudes to age rather than tinnitus. Consistent with our findings, studies that only included young, normal-hearing subjects showed no differences in ABR and EFR between individuals with and without tinnitus (Gilles et al., 2016; Guest et al., 2017).

Nevertheless, other studies did reveal decreases in electrophysiological markers related to tinnitus. Paul et al. (2017) observed reduced EFR responses in subjects with tinnitus, while they reduce age-related effects by only including young adults. A possible explanation for these different findings could be the EFR stimulus characteristics. In comparison to the referenced study, we utilized a higher modulation rate of 110 Hz, which is more reflective of activity closer to the auditory nerve. Lower rates, such as 85 Hz in Paul et al. (2017), are typically associated with midbrain responses. These reduced responses towards more cortical regions were also observed by assessing cortical auditory evoked potentials (Cardon et al., 2022). Additionally, we presented a RAM-stimulus instead of the sinusoidal modulation envelope used in Paul et al. (2017), because RAM-stimuli are less sensitive to OHC-damage and more sensitive to CS (Vasilkov et al., 2021). Schaette and McAlpine (2011) also observed, contrary to our findings, reduced ABR wave-I amplitudes in subjects with tinnitus. This may be explained by the slightly older age of the young groups (around 35 years) or again by potential OHC damage. Previous studies have shown that ABR wave I is also affected by OHC damage, measurable by OAEs (Vasilkov & Verhulst, 2019; Verhulst et al., 2016). However, OAEs were not measured in either study. Future electrophysiological studies should include OAE measurements to examine the effects of OHC loss, since this is typically associated with tinnitus (Degeest et al., 2014).

Gilles et al. (2016) included TEOAE and DPOAE measurements in combination with ABR and did not find any significant tinnitus-related group differences in either measurement. Contrary, Bramhall et al. (2018) found significant tinnitus-related differences in young participants for both DPOAE and 4 kHz tone burst ABR. This ABR wave-I reduction remained after adjusting for DPOAE levels. In a following study that included larger groups with a greater age range, they concluded that both ABR wave I and DPOAEs could be associated with greater probability of tinnitus (Bramhall, McMillan, et al., 2019). EFR measurements were not included, nor were tinnitus or hyperacusis questionnaires collected in that study. Taking this study into account, the frequency specificity of our electrophysiological markers may also explain the lack of tinnitus-related differences. The click ABR represented frequencies between 1000 and 4000 Hz, whereas EFR only represented 4000 Hz. Future research could focus on measuring EFR and tone-burst ABR around the tinnitus pitch.

In addition to the three markers discussed, the middle ear muscle reflex (MEMR) has been suggested as a possible marker for CS (Bharadwaj et al., 2019). Mixed results have been found using this technique in tinnitus studies (Casolani et al., 2022; Guest et al., 2019; Wojtczak et al., 2017). When considering this marker of hidden hearing loss, age and OHC damage should also be considered as possible confounders. Further investigation is needed to determine the link between MEMR and tinnitus, in combination with electrophysiological measurements to prove a relation between CS and tinnitus. A recent tinnitus study by Vasilkov et al. (2023) included both ABR and MEMR measurements, revealing significant tinnitus-related group differences for both markers. However, this study by Vasilkov et al. (2023) covered a wide age range, from 18 to 72 years old, and only considered age based on hearing thresholds, overlooking potential insights related to hidden hearing loss and cochlear synaptopathy within the examined cohorts.

Additionally to the ABR wave-I analysis, central gain was evaluated using the wave-I/V ratio. Although not significant, only age-related group differences were observed for this I/V ratio. This aligns again with the findings of Johannesen and Lopez-Poveda (2021), concluding that ABR-based central gain is associated with age rather than tinnitus. Harris et al. (2022) investigated this age-related central gain effect, identifying hyperexcitability in cortical activity despite reduced AN input. This effect was related to lower cortical GABA levels and poorer SPIN perception, independent from hearing thresholds (Harris et al., 2022). Gu et al. (2012) also addressed the importance of age in their ABR study, noting that older subjects (around 40 years of age) showed reduced ABR wave-I amplitudes compared to a younger group (around 20 years of age). This study revealed a central gain effect at high click intensities of 80 dB nHL, corresponding to 120 dB peSPL, in older subjects with tinnitus. However, young subjects with tinnitus were not included in this study.

### 4.2 TFS coding may be an overlooked factor in tinnitus research

As a functional representation of hidden-hearing-loss complaints, speech (in noise) intelligibility was examined. Besides the observed age-related reduction in HP-filtered speech, older individuals with tinnitus unexpectedly showed significantly improved LP SPIN scores compared to those without tinnitus. Curiously, after exploring possible related factors, these scores did not correlate with hearing thresholds or electrophysiological measures. Instead, a significant correlation was found with tinnitus distress, based on the THI and especially TFI questionnaires. This correlation may imply that individuals with higher tinnitus distress were more focused on, or more sensitive to LP-SPIN information. The HP-filtered SPIN condition did not yield a similar tinnitus-related effect, suggesting that a low-frequency cue would be responsible for these findings. Age-related differences, as observed in the HP condition, can be attributed to several factors, such as higher hearing thresholds, hidden hearing loss, or cognitive decline associated with aging. Explaining these differences is beyond the scope of this study. However, our tinnitus-related difference in LP SPIN scores within the older group cannot be attributed to age-related confounding factors since no significant differences were observed in parameters for hidden hearing loss and audiometric thresholds.

The LP and HP filtering of the speech understanding tests were chosen to distinguish between TFS and TENV processing. It is hypothesized that for LP stimuli, humans utilize both TFS and TENV information, while for HP stimuli, one is restricted to TENV information due to the human phase locking limit at 1500 Hz (Bharadwaj et al., 2014; Brughera et al., 2013; Joris et al., 2004; Verschooten et al., 2019). Based on the age-related reduction that we observed in HP SPIN and EFR measurements, we suggest that older participants had worse TENV encoding. Since this reduction in TENV encoding was observed in both older groups with and without tinnitus, we suggest that our observed tinnitus-related difference could be explained by TFS processing. Given that this improvement in SPIN was only observed in the LP condition, we hypothesize that older individuals with tinnitus may rely more strongly on TFS information than older subjects without tinnitus.

This improved TFS encoding could be related to the findings of Bureš et al. (2019), who reported elevated scores in detecting suprathreshold intensity changes in their tinnitus group, indicating a heightened sensitivity to suprathreshold temporal encoding. Similarly, Zeng et al. (2020) observed enhanced scores in intensity discrimination, while frequency discrimination was poorer in the tinnitus group. Correlations between these psychoacoustic tests and THI-scores were not examined, nor were hyperacusis questionnaires incorporated. Suprathreshold TFS and TENV encoding should be further explored in future tinnitus studies. Moon et al. (2015) already performed TFS and TENV-related psychoacoustic measurements in their study, making interaural comparisons within unilateral and bilateral tinnitus patients. However, their study lacked a non-tinnitus reference group for comparison with our findings.

Although there is a lack of TFS-related research in tinnitus, many studies have already focused on speech understanding. In contrast to our findings, several studies have reported reduced speech understanding in normal hearing patients with tinnitus, particularly in noisy environments (Gilles et al., 2016; Ivansic et al., 2017; Jagoda et al., 2018; Sommerhalder et al., 2023). The variability in results across studies may be due to different speech stimuli, noise conditions, the influence of cognitive factors, and heterogeneity of the tinnitus groups (Ivansic et al., 2017; Tai & Husain, 2019). None of these studies applied filtering to speech stimuli or attempted to address TFS coding. One study did perform a test using LP filtered monosyllabic words to address temporal lobe dysfunction, with a 500 Hz cutoff (Goldstein & Shulman, 1999). However, these stimuli and objective are not comparable with the filtered sentence test at 1500 Hz cutoff that we used to address TFS encoding. Larger sample sizes are needed to further investigate the link between tinnitus distress and improved speech comprehension, which was not the original objective of this study. Future studies should further explore the use of LP and HP filtering on speech in noise tests to confirm our results in a more extensive cohort. Also, the role of attention and other cognitive mechanisms necessary for understanding LP SPIN and related to tinnitus, should be further investigated.

### 4.3 Hyperacusis could explain improved speech in noise scores

Hyperacusis could also explain the improved LP SPIN scores. Despite the limited sample size, the grouping based on the HQ suggests that individuals with hyperacusis tend to score better on the LP SPIN test compared to both the non-tinnitus group and the tinnitus group without hyperacusis. This trend could suggest that individuals with hyperacusis may be particularly sensitive to low-frequency sound, in contrast to the high-frequent reductions in dynamic range that are typically observed in these patients. Auerbach et al. (2014) performed a comprehensive study on neural central gain enhancement and reported that loss of inhibition and enhanced central gain could especially be visible in low frequencies at the colliculus inferior. Since our electrophysiological measurements were based on high-frequent stimuli, further exploration of low frequency encoding could be valuable in tinnitus research.

Since THI, TFI and HQ all seem to be related to improved LP SPIN, it is difficult to attribute the improved LP SPIN scores to either tinnitus distress or hyperacusis. Previous studies on hyperacusis have consistently shown elevated THI and TFI scores, indicating that tinnitus distress and hyperacusis are related to each other (Degeest et al., 2016; Fioretti et al., 2013; Jacquemin et al., 2022). These higher THI and TFI scores were also visible in our older groups with hyperacusis, despite the small group size. As hyperacusis is reported in approximately 40% of tinnitus subjects, more comprehensive studies are needed to distinguish between subgroups with and without tinnitus, and with and without hyperacusis (Ralli et al., 2017).

Recent literature highlights the importance of distinguishing between tinnitus with and without hyperacusis. For example, Refat et al. (2021) showed contrasting results of ABR-amplitudes when comparing tinnitus subjects with and without hyperacusis to a non-tinnitus reference group. Hyperacusis participants exhibited stronger wave-V amplitudes, while participants with only tinnitus showed lower amplitudes. Zeng (2020) proposed different models for hyperacusis and non-hyperacusis, indicating that tinnitus may result from additive central noise due to traditional hearing loss, while hyperacusis occurs with steeper loudness growth due to central gain enhancement caused by hidden hearing loss. Similarly, Chen et al. (2021) proposed a model in which tinnitus is accounted for by central enhancement at low levels, and therefore high-SR fibers, while hyperacusis is related to neural gain at high levels, and thus low-SR fibers. Future research should include tinnitus subjects with or without hyperacusis, as well as hyperacusis subjects without tinnitus. Furthermore, the use of extended hyperacusis questionnaires, such as the Hyperacusis Impact Questionnaire and the Inventory of Hyperacusis Symptoms, would improve the understanding of the interplay between tinnitus and hyperacusis (Aazh et al., 2022; Greenberg & Carlos, 2018).

## 5 Conclusion

This study examined the relationship between tinnitus and potential markers for hidden hearing loss, namely EHF thresholds, ABR wave I amplitude and EFR magnitude. The results showed clear age-related effects in peripheral hidden hearing loss markers, while no significant tinnitus-related differences were observed. Based on these findings, cochlear synaptopathy is unlikely to be the driving force behind tinnitus. Only in a LP SPIN condition, improved scores were observed for the older tinnitus subjects compared to the older group without tinnitus. Based on this exploratory data, we hypothesize that the improved LP SPIN scores in tinnitus patients could be explained by TFS- or hyperacusis-related mechanisms. Based on these findings, we propose the following recommendations for future research:

- When performing electrophysiological measurements on participants with tinnitus, hidden hearing loss due to age should be taken into account. Incorporating age grouping into study designs can eliminate confounding factors such as hidden hearing loss and cochlear synaptopathy.
- Future studies on tinnitus should include measurements for low-frequency encoding, and more specifically TFS coding. A more detailed exploration of TFS coding can contribute significantly to unraveling the mechanisms underlying tinnitus and hyperacusis.
- Besides tinnitus complaints, hyperacusis symptoms should also be assessed thoroughly in future research. This requires grouping based on hyperacusis and tinnitus and including detailed hyperacusis questionnaires.

## 6 Acknowledgement

We thank Emma De Smet and Neri Spitaels for patient recruitment and data-collection as part of their Msc studies in audiology at Ghent University.

## 7 Formatting of funding sources

Work supported by Horizon 2020 ERC-StG 678120 RobSpear; and FWO G068621N AuDiMod.

## 8 Autor contributions

Conceptualization (SV, HK), Methodology (SV, SK, HK), Formal analysis (SK, SV, HK, BT, PD), Writing – original draft (PD), Writing – review & editing (HK, ID, SV), Funding acquisition (SV)

## Abbreviations

ABR: Auditory Brainstem Response
ANFs: Afferent Nerve Fibers
BB: Broadband
CS: Cochlear Synaptopathy
DPOAE: Distortion Product Otoacoustic Emission
EEG: Electroencephalography
EFR: Envelope Following Response
EHF: Extended High Frequency
HP: High Pass
HQ: Hyperacusis Questionnaire
LP: Low Pass
MEMR: Middle Ear Muscle Reflex
SRT: Speech Reception Threshold
SR: Spike Rate
SPIN: Speech Perception In Noise
SPIQ: Speech Perception In Quiet
TENV: Temporal Envelope
THI: Tinnitus Handicap Inventory
TFI: Tinnitus Functional Index
TFS: Temporal Fine Structure
VAS: Visual Analogue Scale

